# ViralEpiBase: a manually curated repository of epitranscriptomic modification sites across viral RNA genomes and virus-encoded transcripts

**DOI:** 10.64898/2026.07.02.735974

**Authors:** Sreeram Srinivasan, Ajit Chande

**Affiliations:** From the Molecular Virology Laboratory, Department of Biological Sciences, Indian Institute of Science Education and Research (IISER) Bhopal, Bhopal 462066, Madhya Pradesh, India

## Abstract

Post-transcriptional chemical modifications of RNA, collectively termed the epitranscriptome, have emerged as critical regulatory layers governing viral replication, pathogenicity, and host– virus interactions. Despite the rapid accumulation of experimental data on viral RNA modifications, no dedicated, freely accessible resource existed for systematically cataloguing these sites across diverse viral species. Here we present ViralEpiBase, a manually curated database of epitranscriptomic modification sites identified in viral RNA genomes and virus-encoded transcripts at single-nucleotide resolution. ViralEpiBase currently integrates seven chemically distinct RNA modification types: N6-methyladenosine (m6A), N1-methyladenosine (m1A), pseudouridine (Ψ), 5-methylcytosine (m5C), 2′-O-methylation (2′OMe), inosine and N4-acetylcytidine (ac4C); across 12 viral species encompassing both DNA and RNA viruses of clinical and biological significance. Each entry is linked to its primary literature source or deposited dataset and is retrievable by modification type, genomic coordinates, or viral taxonomy. The database is freely accessible through an intuitive web interface and is updated continuously as new experimental evidence becomes available. ViralEpiBase thus provides the first unified platform dedicated exclusively to viral epitranscriptomics and is designed to facilitate mechanistic investigation of RNA modification functions in viral biology.

## Introduction

Viruses remain a principal cause of human morbidity and mortality, with emerging and re-emerging pathogens continuously challenging global public health infrastructures (1). Central to viral pathogenesis is their capacity to subvert host cellular machinery by utilizing them for their replication, transcription, translation, and for immune evasion(2,3). RNA molecules encoded by viruses are subject to diverse regulatory mechanisms, among which post-transcriptional chemical modifications of ribonucleotides have attracted increasing attention as determinants of viral RNA fate, stability, and translatability(4).

The epitranscriptome comprises over 170 chemically distinct RNA modifications, most of which have been identified in cellular RNAs, with only a subset has been characterized in viral RNA(4,5). Among these, N6-methyladenosine (m6A) is the most abundantly studied and has been demonstrated to modulate the replication cycles and immune evasion strategies of diverse viruses, including Hepatitis B virus, Flaviviridae members, and retroviruses (6–10). Additional modification types including m1A, pseudouridine, m5C, 2′-O-methylation, inosine, and ac4C are increasingly recognized as regulatory features of both host and viral RNA(4). These modifications collectively influence viral RNA secondary structure, protein–RNA interactions, innate immune sensing, and translational efficiency, thereby constituting a layer of epitranscriptomic regulation with direct consequences for viral infectivity and pathogenesis(4,10,11).

Bioinformatic resources for exploring RNA modifications have expanded considerably over the past decade, which includesdatabases like RMBase v3.0 (12), MODOMICS(13), Sci-ModoM(14), POSTAR3(15), and RMVar 2.0(16) provide broad coverage of modifications across mammalian and other eukaryotic transcriptomes. However, these repositories focus predominantly on cellular organisms and afford only limited coverage of viral systems. The sole existing resource with a viral dimension, DirectRMDB(17), which is restricted only to Chikungunya virus and Influenza A virus, thereby creating a significant gap to support the growing community of researchers investigating viral epitranscriptomics.

To address this unmet need, we developed ViralEpiBase, a manually curated, freely accessible database dedicated exclusively to viral epitranscriptomics. ViralEpiBase integrates experimentally validated RNA modification sites from peer-reviewed literature and public repositories across 12 viral species, spanning both DNA and RNA viruses. By providing positional-resolution data on seven modification types alongside links to sources, ViralEpiBase enables systematic exploration of the viral epitranscriptome and is intended to serve as a community resource facilitating mechanistic and translational research on RNA modifications in viral biology.

## Materials and Methods

### Data curation strategy

A systematic literature search was conducted using PubMed to identify peer-reviewed studies reporting post-transcriptional RNA modifications in viral genomes or virus-encoded transcripts. Search terms included combinations of modification-type keywords (e.g., “m6A”, “pseudouridine”, “m5C”, “RNA methylation”, “inosine “) with viral taxonomy terms. High-throughput modification datasets deposited in [NCBI-GEO](https://www.ncbi.nlm.nih.gov/geo/query/acc.cgi) were additionally retrieved using cognate accession identifiers cited in curated publications (Supplementary File S1). For Chikungunya virus and Influenza A virus, modification data from DirectRMDB(17), were incorporated. All retrieved records underwent manual inspection for verifying the quality, modification type, chromosomal coordinates, and the identity of the viral RNA element (genomic RNA vs. virus-encoded transcript).

### Database architecture and web interface

The ViralEpiBase web application employs a three-tier architecture. The frontend interface is constructed using HTML, CSS, and JavaScript, providing a responsive and browser-compatible user experience. The server-side application logic is implemented in Python using the Flask micro-framework, which handles routing, query processing, and data serialisation. All curated modification records are stored in a MongoDB database, organised within a single collection with documents structured according to a standardised schema encompassing viral species, modification type, genomic coordinate, strand, dataset source. MongoDB’s document-oriented model affords flexible indexing strategies enabling rapid retrieval across heterogeneous query types. The application is deployed on a Linux server accessible at https://viralepibase.iiserb.ac.in.

### Data availability and download

All curated data are available for bulk download from the Downloads section of the website. The database will be maintained and updated with the availability of new published experimentally validated modification data for viral species. In accordance with the availability policy of this journal, ViralEpiBase will remain freely and openly accessible for a minimum of two years following publication, with ongoing maintenance supported through institutional resources at IISER Bhopal.

## Results

### Overview of the ViralEpiBase

ViralEpiBase currently catalogues experimentally validated RNA modification sites across 12 viral species, spanning seven chemically distinct modification types: m6A, m1A, pseudouridine (Ψ), m5C, 2′-O-methylation (2′OMe), inosine (A-to-I editing), and ac4C (Table 1). The curated viral species comprises both DNA viruses (e.g., members of the Herpesviridae and Papillomaviridae families) and RNA viruses (e.g., Flaviviridae, Retroviridae, and Orthomyxoviridae), reflecting the taxonomic breadth of current viral epitranscriptomic literature. Modification sites are annotated at single-nucleotide resolution wherever the source experimental data permit, including genomic coordinates referenced against standard RefSeq viral genome assemblies, strand information, identity of the modified RNA element (viral genomic RNA, or virus encoded transcript and the source from which the information was curated.

**Table 1.** Summary of RNA modification types and viral species represented in ViralEpiBase.

| Modification type | Chemical abbreviation | Viral species represented (examples) |
| --- | --- | --- |
| N6-methyladenosine | m6A | HIV-1, HBV, HCV, DENV, ZIKV, IAV, HPV, EBV, KSHV |
| N1-methyladenosine | m1A | HIV-1, IAV |
| Pseudouridine | ψ | HIV-1, IAV, EBV |
| 5-methylcytosine | m5C | HIV-1, CHIKV, IAV, SARS-CoV-2 |
| 2'-O-methylation | 2'OMe | HIV-1, HCV, IAV |
| Inosine (A-to-I) | I | HIV-1, EBV, IAV |
| N4-acetylcytidine | ac4C | HIV-1, IAV |
| Total across all types | — | 12 viral species |

### Web interface and user query functionality

The ViralEpiBase web interface is structured around five primary sections: Home, DNA Virus, RNA Virus, Help, and Downloads (Figure-1). The [Home page](https://viralepibase.iiserb.ac.in/#homeSection) provides an introductory overview of the database, its scope, and summary statistics for the current release. The [DNA Virus](https://viralepibase.iiserb.ac.in/#dnaVirusSection) and [RNA Virus](https://viralepibase.iiserb.ac.in/#rnaVirusSection) sections each provide interactive query interfaces that allow users to retrieve modification records on selecting the virus of interest by (i) modification type (e.g., selecting m5C returns all m5C sites in that virus), (ii) genomic coordinates or position range. Query results are displayed in a tabular format presenting the modification type, genomic position, strand, RNA element, detection method, experimental context, and the primary literature citation or GEO accession number. The [Help section](https://viralepibase.iiserb.ac.in/#helpSection) contains a step-by-step tutorial describing each query mode with annotated screenshots (Supplementary File S2). The [Download section](https://viralepibase.iiserb.ac.in/#downloadSection) provides unrestricted download of the full database.

**Figure 1.**
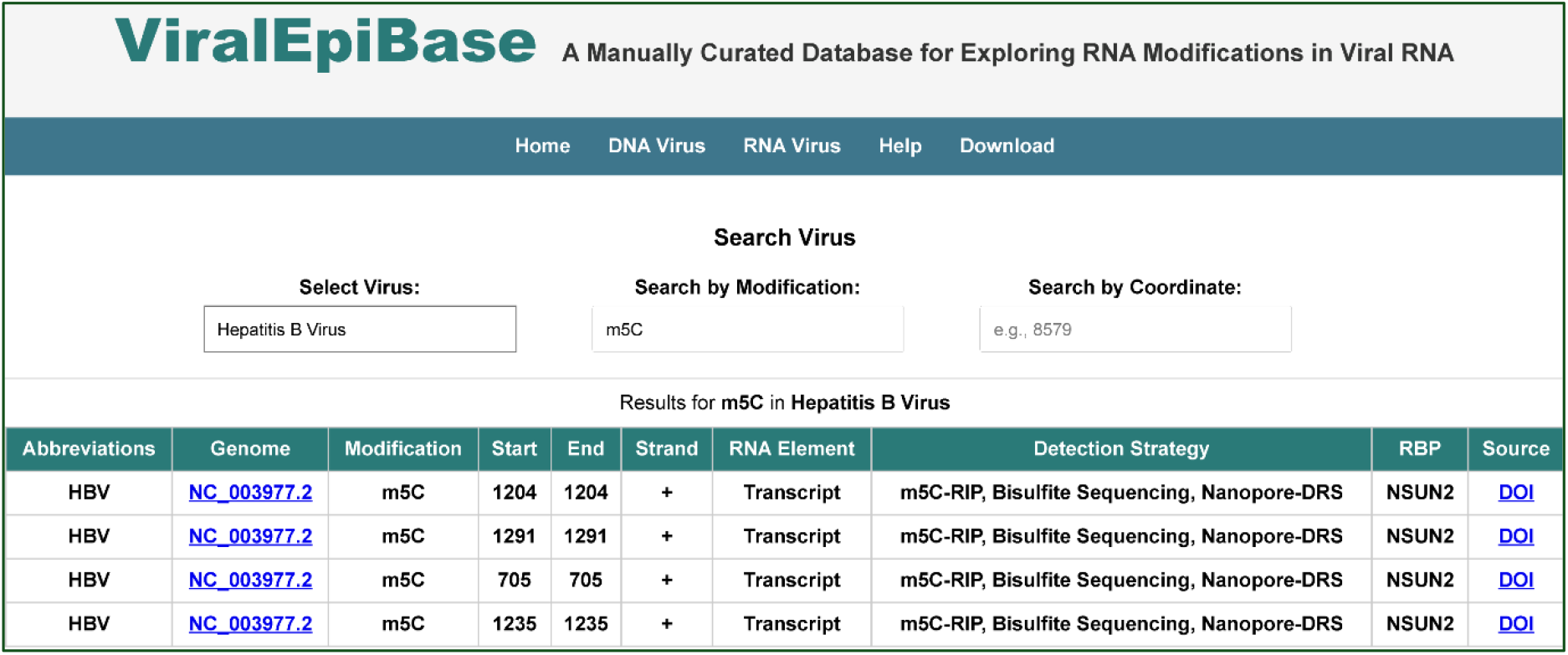
Overview of the ViralEpiBase web interface. Home page showing database statistics and navigation structure. Representative query result table for m5C modifications, showing position, RNA Element, Detection Strategy, RBP mediating the modification and source reference fields.

## Discussion

Epitranscriptomics has transformed our understanding of gene regulation, revealing that reversible chemical modifications of RNA represent a dynamic layer of post-transcriptional control analogous to epigenetic regulation of DNA(4,6). In the context of virology, epitranscriptomic modifications have been implicated in nearly every stage of the viral replication cycle. m6A modifications on viral RNA have been shown to modulate cap-independent translation, facilitate nuclear export of viral transcripts, suppress innate immune sensing, and promote or antagonise viral replication depending on the biological context(7–9,11). Similarly, m5C modifications have been documented in the genomes of HIV-1 and SARS-CoV-2, where they may influence RNA stability and their processing (18,19). A-to-I editing by ADAR enzymes affects the genomes of multiple RNA viruses, with consequences for viral protein diversity and immune evasion(20). Despite this rapidly expanding literature, no unified computational resource existed to consolidate these observations in a searchable, structured form, a gap that ViralEpiBase directly fills.

Existing databases for RNA modifications revolve around cellular transcriptomes and do not cover viruses. RMBase v3.0(12) and MODOMICS(13) provide comprehensive coverage of eukaryotic modification sites but doesnt prioritises viral RNA as a distinct data category. Sci-ModoM(14) ntegrates high-throughput stoichiometric data across the human transcriptome but does not accommodate viral RNA. DirectRMDB (17), the most closely related resource, compiles modification sites detected by direct nanopore RNA sequencing technology but is currently restricted to two viral species (Chikungunya virus and Influenza A virus), and its scope is defined by the detection technology rather than the biological system. ViralEpiBase extends this coverage to 12 viral species and seven modification types, integrating data derived from multiple experimental platforms including MeRIP-seq, m6A-crosslinking and immunoprecipitation (m6A-CLIP), bisulfite sequencing, and direct RNA nanopore sequencing. This heterogeneous evidence base reflects the current state of the viral epitranscriptomics literature and is an inherent feature of a manually curated, literature-driven resource.

A key strength of ViralEpiBase is its commitment to manual curation. The inclusion of source metadata for every record — detection method, experimental condition, primary citation, and GEO accession where applicable allows users to critically evaluate the evidence quality of individual entries and to stratify analyses by detection platform. We acknowledge that manual curation introduces inherent limitations of scale and completeness; as the viral epitranscriptomics literature continues to expand, semi-automated curation workflows integrating natural language processing and machine learning may be incorporated in future releases to support data throughput without compromising accuracy.

Future development of ViralEpiBase will focus on expanding additional viral species and modification types as published data become available, with particular attention to emerging pathogens. Secondly, functional annotation layers will be integrated, i.e., protein–RNA interactions influenced by specific modification sites, and phenotypic consequences of modification in the context of viral replication. Third, interoperability with existing viral genomics and proteomics resources including [ViPR](http://www.viprbrc.org/)(21), [ICTV](https://ictv.global/)(22), and [UniProt](https://www.uniprot.org/)(23) will be implemented through standardised data exchange formats to enhance cross-database querying capacity. These enhancements are expected to transform ViralEpiBase from an evidence repository into a hypothesis-generating platform for the viral epitranscriptomics community.

## Conclusion

ViralEpiBase represents the first dedicated, manually curated database for viral epitranscriptomics, providing positional-resolution annotations of seven RNA modification types across 12 viral species. By consolidating experimentally validated modification sites within a freely accessible, searchable web platform, ViralEpiBase addresses a long-standing gap in the landscape of bioinformatic resources available to virologists and RNA biologists. The database is expected to serve as an enabling resource for mechanistic studies of RNA modification functions in viral replication and pathogenicity, for comparative epitranscriptomics across viral taxonomies, and for the identification of novel therapeutic targets embedded within the viral epitranscriptome. Continued curation and functional annotation of ViralEpiBase will track the growth of this rapidly evolving field.

## Supplementary Data

Supplementary data are available at Database Online.

**Supplementary File S1**. List of all Modifications across Viral RNAssources ubyManual Curation from PubMed and NCBI GEO.

**Supplementary File S2**. ViralEpiBase Tutorial PDF.

## Acknowledgements

The authors thank all members of the Molecular Virology Laboratory at IISER Bhopal for productive discussions. R. Dalavi and A. Singh are specifically acknowledged for their technical assistance and critical evaluation of the database interface.

## Author Contributions

Conceptualization and supervision: A.C. Data curation: S.S. Web interface design and implementation: S.S. Original manuscript preparation: S.S. Critical manuscript review and editing: A.C. Funding acquisition: A.C.

## References

1. Evolving viral threats. Nature Reviews Microbiology. Nature Research; 2026. p. 1–2. doi:10.1038/s41579-025-01263-x PubMed PMID: 41381702.

2. Bhardwaj V, Singh A, Choudhary A, Dalavi R, Ralte L, Chawngthu RL, et al. HIV-1 Vpr induces ciTRAN to prevent transcriptional repression of the provirus [Internet]. 2023. Available from: https://www.science.org

3. Iselin L, Palmalux N, Kamel W, Simmonds P, Mohammed S, Castello A. Uncovering viral RNA–host cell interactions on a proteome-wide scale. Trends in Biochemical Sciences. Elsevier Ltd; 2022. p. 23–38. doi:10.1016/j.tibs.2021.08.002 PubMed PMID: 34509361.

4. Tian Y, Wang X. RNA modification is the mark and strategy for host-microbe interactions. Cellular and Molecular Life Sciences. Springer Science and Business Media Deutschland GmbH; 2025. doi:10.1007/s00018-025-05842-2 PubMed PMID: 40779243.

5. Wei J, He C. RNA modifications in gene regulation: Functions and pathways. Cell. Elsevier B.V.; 2026. p. 1591–619. doi:10.1016/j.cell.2026.01.006

6. Imam H, Khan M, Gokhale NS, McIntyre ABR, Kim GW, Jang JY, et al. N6-methyladenosine modification of hepatitis b virus RNA differentially regulates the viral life cycle. Proc Natl Acad Sci U S A. 2018 Aug 28;115(35):8829–34. doi:10.1073/pnas.1808319115 PubMed PMID: 30104368.

7. Zhao J, Lee EE, Kim J, Yang R, Chamseddin B, Ni C, et al. Transforming activity of an oncoprotein-encoding circular RNA from human papillomavirus. Nat Commun. 2019 Dec 1;10(1). doi:10.1038/s41467-019-10246-5 PubMed PMID: 31127091.

8. Gokhale NS, McIntyre ABR, McFadden MJ, Roder AE, Kennedy EM, Gandara JA, et al. N6-Methyladenosine in Flaviviridae Viral RNA Genomes Regulates Infection. Cell Host Microbe. 2016 Nov 9;20(5):654–65. doi:10.1016/j.chom.2016.09.015 PubMed PMID: 27773535.

9. Gonzales-van Horn SR, Sarnow P. Making the Mark: The Role of Adenosine Modifications in the Life Cycle of RNA Viruses. Cell Host and Microbe. Cell Press; 2017. p. 661–9. doi:10.1016/j.chom.2017.05.008 PubMed PMID: 28618265.

10. Aufgebauer CJ, Bland KM, Horner SM. Modifying the antiviral innate immune response by selective writing, erasing, and reading of m6A on viral and cellular RNA. Cell Chemical Biology. Elsevier Ltd; 2024. p. 100–9. doi:10.1016/j.chembiol.2023.12.004 PubMed PMID: 38176419.

11. Ding S, Liu H, Liu L, Ma L, Chen Z, Zhu M, et al. Epigenetic addition of m5C to HBV transcripts promotes viral replication and evasion of innate antiviral responses. Cell Death Dis. 2024 Jan 1;15(1). doi:10.1038/s41419-023-06412-9 PubMed PMID: 38216565.

12. Xuan J, Chen L, Chen Z, Pang J, Huang J, Lin J, et al. RMBase v3.0: decode the landscape, mechanismsãnd functions of RNA modifications. Nucleic Acids Res. 2024 Jan 5;52(D1):D273–84. doi:10.1093/nar/gkad1070 PubMed PMID: 37956310.

13. Sordyl D, Boileau E, Bernat A, Maiti S, Mukherjee S, Moafinejad SN, et al. MODOMICS: a database of RNA modifications and related information. 2025 update and 20th anniversary. Nucleic Acids Res. 2026 Jan 6;54(D1):D219–25. doi:10.1093/nar/gkaf1284 PubMed PMID: 41277531.

14. Boileau E, Wilhelmi H, Busch A, Cappannini A, Hildebrand A, Bujnicki JM, et al. Sci-ModoM: a quantitative database of transcriptome-wide high-throughput RNA modification sites. Nucleic Acids Res. 2025 Jan 6;53(D1):D310–7. doi:10.1093/nar/gkae972 PubMed PMID: 39498498.

15. Zhao W, Zhang S, Zhu Y, Xi X, Bao P, Ma Z, et al. POSTAR3: An updated platform for exploring post-transcriptional regulation coordinated by RNA-binding proteins. Nucleic Acids Res. 2022 Jan 7;50(D1):D287–94. doi:10.1093/nar/gkab702 PubMed PMID: 34403477.

16. Huang Y, Zhang L, Mu W, Zheng M, Bao X, Li H, et al. RMVar 2.0: an updated database of functional variants in RNA modifications. Nucleic Acids Res. 2025 Jan 6;53(D1):D275–83. doi:10.1093/nar/gkae924

17. Zhang Y, Jiang J, Ma J, Wei Z, Wang Y, Song B, et al. DirectRMDB: a database of post-transcriptional RNA modifications unveiled from direct RNA sequencing technology. Nucleic Acids Res. 2023 Jan 6;51(D1):D106–16. doi:10.1093/nar/gkac1061 PubMed PMID: 36382409.

18. Wang H, Feng J, Fu Z, Xu T, Liu J, Yang S, et al. Epitranscriptomic m 5 C methylation of SARS-CoV-2 RNA regulates viral replication and the virulence of progeny viruses in the new infection. Sci. Adv [Internet]. 2024. Available from: https://www.science.org

19. Courtney DG, Tsai K, Bogerd HP, Kennedy EM, Law BA, Emery A, et al. Epitranscriptomic Addition of m5C to HIV-1 Transcripts Regulates Viral Gene Expression. Cell Host Microbe. 2019 Aug 14;26(2):217-227.e6. doi:10.1016/j.chom.2019.07.005 PubMed PMID: 31415754.

20. Piontkivska H, Wales-McGrath B, Miyamoto M, Wayne ML. ADAR Editing in Viruses: An Evolutionary Force to Reckon with. Genome Biol Evol. 2021 Nov 1;13(11). doi:10.1093/gbe/evab240 PubMed PMID: 34694399.

21. Pickett BE, Sadat EL, Zhang Y, Noronha JM, Squires RB, Hunt V, et al. ViPR: An open bioinformatics database and analysis resource for virology research. Nucleic Acids Res. 2012 Jan;40(D1). doi:10.1093/nar/gkr859 PubMed PMID: 22006842.

22. Black EJ, Powell CS, Dempsey DM, Hendrickson RC, Mims LR, Lefkowitz EJ. Virus taxonomy: the database of the International Committee on Taxonomy of Viruses. Nucleic Acids Res. 2026 Jan 6;54(D1):D776–89. doi:10.1093/nar/gkaf1159

23. Bateman A, Martin MJ, Orchard S, Magrane M, Adesina A, Ahmad S, et al. UniProt: the Universal Protein Knowledgebase in 2025. Nucleic Acids Res. 2025 Jan 6;53(D1):D609–17. doi:10.1093/nar/gkae1010 PubMed PMID: 39552041.

